# Nano-positioning and tubuline conformation determine transport of mitochondria along microtubules

**DOI:** 10.1101/2020.04.27.064766

**Authors:** V. Van Steenbergen, F. Lavoie-Cardinal, Y. Kazwiny, M. Decet, T. Martens, P. Verstreken, W. Boesmans, P. De Koninck, P. Vanden Berghe

## Abstract

Correct spatiotemporal distribution of organelles and vesicles is crucial for healthy cell functioning and is regulated by intracellular transport mechanisms. Controlled transport of bulky mitochondria is especially important in polarized cells such as neurons that rely on these organelles to locally produce energy and buffer calcium. Mitochondrial transport requires and depends on microtubules which fill much of the available axonal space. How mitochondrial transport is affected by their position within the microtubule bundles is not known. Here, we found that anterograde transport, driven by kinesin motors, is susceptible to the molecular conformation of tubulin both *in vitro* and *in vivo*. Anterograde velocities negatively correlate with the density of elongated tubulin dimers, similar to GTP-tubulin, that are more straight and rigid. The impact of the tubulin conformation depends primarily on where a mitochondrion is positioned, either within or at the rim of microtubule bundle. Increasing elongated tubulin levels lowers the number of motile anterograde mitochondria within the microtubule bundle and increases anterograde transport speed at the microtubule bundle rim. We demonstrate that the increased kinesin step processivity on microtubules consisting of elongated dimers underlies increased mitochondrial dynamics. Our work indicates that the molecular conformation of tubulin controls mitochondrial motility and as such locally regulates the distribution of mitochondria along axons.

## Introduction

Mitochondria are found in almost all eukaryotic cell types and are of key importance to maintain intracellular processes such as adenosine triphosphate (ATP) production, calcium homeostasis, stress responses and cell fate ^1-3^. As mitochondria are formed in the cell soma, active transport towards sites with a high energy demand is required for healthy cellular function. The long, narrow and highly branched axonal and dendritic projections of nerve cells present an extra challenge for transport of these organelles to their target sites and require precise spatiotemporal regulation of the transport machinery ^4-7^. Inefficient organization of the molecular machinery can lead to mitochondrial transport deficits, which are implicated in neurodegenerative diseases such as Amyotrophic lateral sclerosis (ALS), Alzheimer’s and Huntington’s disease ^8-10^.

The molecular railroads used for long-distance intracellular transport are microtubules, long tubulin polymers that provide binding sites for kinesin and dynein motor proteins ^11^. Upon polymerization, tubulin dimers bind in an exchangeable manner to guanosine triphosphate (GTP) at the ß-tubulin subunit, favoring a stable, elongated dimer conformation. Once incorporated in the tubular lattice, GTP hydrolysis leads to a conformational change towards a less stable and compacted guanosine diphosphate (GDP)-bound tubulin dimer. Because GTP hydrolysis lags behind microtubule growth, it was thought that these stable GTP-tubulin dimers were only present at the microtubule growing tip to prevent depolymerization ^12-14^. However, recent studies indicate that GTP-bound tubulin dimers are also present further along the microtubule lattice ^15,16^. These so-called GTP-tubulin islands could serve as rescue sites to prevent depolymerization of the microtubule or provide a location for self-repair ^16-20^.

While it is likely that a high GTP-to GDP-tubulin ratio is required to strengthen the complex architecture in neuronal projections, it also influences binding of kinesin motor proteins. As these motors have been shown to regulate sorting of cargo to the somatodendritic and axonal domains, the presence of GTP-bound tubulin dimers could, apart from influencing general intracellular transport, also be involved in regulating polarized transport ^21-23^. How kinesin binding affinity is regulated by the conformation of tubulin depends on the kinesin subtype. KIF1S binds more weakly to GTP-bound than to GDP-bound tubulin dimers while KIF5 has increased binding affinity to GTP-bound tubulin dimers ^21,24,25^. Moreover, the processive motion of kinesin motors depends on the conformational state of the tubulin dimer, whereby the GTP-bound conformation enables kinesin to move faster on *in vitro* polymerized microtubules ^26^. Recently, T. Shima *et al.* have shown that KIF5C not only preferentially binds GTP-bound tubulin dimers but that it can even pull the conformational state of tubulin from a GDP- to GTP-like state ^25^. However, these findings conflict with effects of drugs that interfere with conformational changes in tubulin dimers such as taxol, a commonly used chemotherapeutic drug. Taxol blocks spindle formation and thus cell division by locking tubulin dimers in an elongated GTP-like tubulin conformation ^27-29^. As taxol raises the amount of stable and elongated tubulin dimers, increased kinesin binding and thus more efficient anterograde transport is expected. However, a conundrum arises as one of taxol’s common side effects is rather an interruption of axonal transport, possibly leading to chemotherapy induced peripheral neuropathy (CIPN) ^30-32^. Thus, how the conformational state of tubulin regulates organelle transport is poorly understood.

In this study, we elucidate how changes in the tubulin dimer conformation locally regulate mitochondrial transport. We show that the presence of elongated tubulin dimers halts anterograde transport specifically of the mitochondria located inside the microtubule bundle, and that this is caused by decreased microtubule flexibility. Consistently, this does not hinder mitochondrial transport that occurs along the rim of the microtubule bundle. On the contrary, we show that this remaining motile fraction is transported faster because of increased processive motion of KIF5B, a kinesin motor for mitochondria that preferentially binds tubulin in an elongated conformation. Finally, using *Drosophila melanogaster* as a model, we show that also increasing the elongated tubulin dimer conformation *in vivo* leads to a reduction in mitochondrial transport in the anterograde direction while transport velocities of the remaining motile fraction are increased.

## Materials and methods

### Primary neuronal cultures

All procedures were approved by the Animal Ethics Committee of the University of Leuven (Belgium) and Université Laval (Canada). All cell cultures apart from these used for live STED imaging were derived from mouse hippocampal tissue. Postnatal day 0-5 C57Bl/6J moue pups were quickly decapitated before dissection. Hippocampi were dissected in sylgard dishes containing cold sterile Hank’s Buffered Salt Solution (HBSS in mM: 5.33 KCl, 0.44 KH_2_PO_4_, 137.93 NaCl, 0.34Na_2_HPO_4_.7H_2_O, 5.56 D-glucose and 10 HEPES). The tissue was minced and incubated in 0.25% trypsin-EDTA (Gibco) supplemented with 80 U/ml DNAse (Roche) for 10 min at 37°C. After three consecutive wash steps with HBSS supplemented with 10% fetal bovine serum (FBS, Sigma Aldrich), the tissue was mechanically dissociated by trituration with syringes with decreasing diameter. Cells were plated at 5 × 10^5^ cells per coverslip (18 mm diameter, coated with poly-D-Lysine) and grown in a 37°C, 5% CO_2_ incubator in Neurobasal-A media (Thermo Fisher Scientific) supplemented with 0.5% penicillin / streptomycin (Lonza), 0.5% B27 (Gibco), 100 ng/ml nerve growth factor (Alomone Labs) and 2mM Glutamax (Thermo Fisher Scientific). Media was replaced 1:1 every three days and cells were used at 7 DIV unless stated otherwise. For live STED recordings, rat hippocampal neurons were prepared as described previously ^33^.

### Transfection

After 6 DIV, expression of TOM20-mCherry-LOVpep and Kif5b-GFP-ePDZb1 (gift from L.C. Kapitein ^34^) was induced in neuronal cultures to guide anterograde transport in axons upon light-induced heterodimerization. Per well, 0.5 µg plasmid was mixed with 0.02% Lipofectamine 2000 reagent (Invitrogen) in Neurobasal-A media and incubated at room temperature for 30 min. The mixture was added dropwise to the wells and media were replaced after 4 hours. Expression was verified the following day.

### Pharmacological treatment

Taxol-treated cultures were incubated with 10 nM taxol dissolved in DMSO (4h, Paclitaxel Cytoskeleton Inc.) in plating medium. Control cultures were treated with an equal amount of DMSO (0.1 %). During imaging, plating medium with DMSO or taxol was replaced by HEPES buffer (in mM: 148 NaCl, 5 KCl, 1 MgCl_2_, 10 Glucose, 10 HEPES, 2 CaCl_2_).

### Mitochondrial transport imaging & analysis

B27 media was replaced by HEPES buffer (in mM: 148 NaCl, 5 KCl, 1 MgCl_2_, 10 Glucose, 10 HEPES, 2 CaCl_2_) for live cell imaging at 37°C. A Zeiss LSM 780 confocal laser scanning microscope (Zeiss) fitted with an Argon laser (488 nm) and solid state lasers (561, 633 nm) was used for mitochondrial imaging (Mitotracker red, 75 nM, 10 min incubation, Thermo Fisher Scientific) in combination with an LD LCI Plan-Apochromat 25x/0.8 Imm Corr DIC M27 water-immersion objective. In-house Igor pro (Wavemetrics, OR, USA) code was used to generate kymographs and analyze mitochondrial transport time lapse recordings as described previously ^35^.

### Electron flow assay isolated mitochondria

Mitochondria were isolated from hippocampal brain tissue using the microTOM22 beads technology (Miltenyi Biotec). The mouse mitochondrial extraction and isolation kit were used for tissue homogenization, digestion and mitochondrial isolation. The isolated mitochondria were then incubated with either taxol in DMSO (10 nM, 1 h) or an equal amount of DMSO as control. These mitochondria were used for the electron flow assay according to the company’s protocol (Agilent Seahorse). In short, mitochondria were resuspended in 9:1 MAS-MSHE buffer and 2 µg mitochondrial protein was plated. The following toxins and substrates were dissolved in MAS buffer before injection: 20 µM rotenone, 100 mM succinate, 40 µM antimycin A and 100 mM ascorbate with 1mM TMPD. Injection volumes were adjusted to ensure a 10 times dilution as a final concentration.

### Immunofluorescence labeling

Cells were fixed with 4% paraformaldehyde (PFA, 30 min, RT) and washed in phosphate buffered saline (PBS). After fixation, cells were incubated in blocking medium containing PBS, 4% serum from secondary hosts (Chemicon International) and 0.1% Triton X-100 (Sigma) (2 h, RT), followed by overnight incubation at 4°C with several primary antibody combinations. Primary antibodies used were: goat TAU (1:1000, cat. number SC1995, Santa Cruz Biotechnology), chicken MAP2 (1:5000, cat. number ab5392, Abcam), rabbit α-tubulin (1:1000, cat. number ab18251, Abcam), rat α-tubulin (1:50, cat. number MA180189, Invitrogen), chicken beta III tubulin (1:50, cat. number ab41489, Abcam), goat kif5b (1:500, cat. Number MBS420641, MyBioSource) and rabbit TOM20 (1:1000, cat. number ab186735, Abcam). After washing with PBS the secondary antibodies were applied (1h, RT). For STED microscopy the following secondary antibodies were used: rabbit-Alexa594 (Invitrogen, 561 excitation) chicken-STAR635P (Abberior, 640 nm excitation), rat-Alexa488 (Invitrogen, 488 excitation), goat-STARRED (Abberior, 640 nm excitation). All antibodies were diluted in blocking medium. Three 10 min wash steps with PBS were performed and excess PBS was removed. All preparations were mounted in Citifluor (Citifluor Ltd.) or Mowiol (Merck) before imaging. For GTP-tubulin staining, we followed the protocol of Dimitrov *et al*. ^15^. Briefly, cells were treated with 0.05% Triton in GPEM buffer (3 min, 37°C) before incubation with MB11 (1:250, cat. number AG-27B-0009-C100 Adipogen) diluted in GPEM buffer supplemented with 2% BSA (15 min, 37°C). After a quick wash in GPEM buffer, donkey anti human alexa 594 (1:1000, cat. number 709-585-149, Jackson ImmunoResearch) was applied (15 min, 37°C) followed by methanol fixation. A Zeiss LSM 780 confocal laser scanning microscope (Zeiss) was used to record fluorescence images.

### Electron microscopy and quantification of microtubule straightness

For electron microscopy, cells were grown on glass coverslips, and fixed with 2.5 % glutaraldehyde in 0.1 M cacodylate buffer (2 h, RT), followed by extensive washing of the samples in the same buffer. Then, the samples were osmicated in 2% osmiumtetroxide in 0.1 M cacodylate buffer (1 h, RT), washed and dehydrated in a graded ethanol series (5 min steps). *En bloc* staining with 3% uranylacetate took place while washing with 70% ethanol (30 min on ice). After washing with 100% ethanol, samples were placed in 1:1 ethanol:epoxy (30 min, on ice) and 1:2 ethanol:epoxy (O/N, RT) mixtures. The next day samples were pre-embedded in a very thin layer of epoxy. Then, regions of interest were selected that were covered with an epoxy filled inverted BEEM-capsule. Finally, samples were cured at 60°C for 2 days. Glass coverslips were removed with hydrofluoric acid (30 min, RT) and washed twice with water. 70 nm sections were cut and post stained with uranyl acetate and lead citrate before imaging with a JEM1400 transmission electron microscope (JEOL) equipped with an EMSIS Quemesa camera (11Mpxl) at 80 kV. To calculate microtubule straightness, the Dendrite straightness tool of Imaris (Bitplane) was used to calculate the deviation of the manually outlined microtubule structure over the weighted general begin to endpoint axes of the microtubule. Microtubules were semi-automatically outlined by two blinded researchers before calculation of microtubule straightness and orientation in XY.

### Stimulated emission depletion (STED) microscopy & colocalisation analysis

An Abberior Expert-Line STED (Abberior Instruments) microscope consisting of an inverted Olympus IX83 microscope body fitted with four pulsed (40 MHz) excitation laser modules (405, 485, 561 and 640 nm), two depletion beams at 595 and 775 nm, a motorized stage with P-736 PINano (Physik Instrumente) and the IX3-ZDC-12 z-drift compensation unit (Olympus) was used for multicolor STED imaging. To relocate cells a 20x Olympus Plan N 0.4 NA air objective was used while a 100x Olympus UplanSApo 1.4 NA oil-immersion objective was used for STED recordings. Emission was detected using a spectral detection module and four avalanche photodiode detectors. For live-cell STED imaging, tubulin was stained with the far-red emitting dye Silicon rhodamine (SiR) using the SiR-tubulin Kit (0.5 μM, 10 min incubation, CY-SC002, Spirochrome) and imaged using 640 nm excitation and 775 nm depletion. Mitochondria were stained with Mitotracker green (75 nM, 10 min incubation, Thermo Fisher Scientific) and imaged in confocal mode using 488 nm excitation. Two-color STED imaging on fixed samples was performed using 775 nm depletion, three-color STED imaging using 775 nm and 595 nm depletion sequentially. In the 2D live-cell imaging configuration, we used 25 × 25 nm pixels, while 40 × 40 × 50 nm pixels were used for fixed 3D STED imaging. To calculate interface percentages between mitochondria and microtubules, Imaris (Bitplane) surface rendering and the surface-to-surface colocalisation Xtension (Bitplane, Matthew Gastinger) were used. To quantify the amount of kinesin dots in proximity of the mitochondrial surface area or interface area between mitochondria and tubulin, kinesin dots were first localized using the ‘spot detection tool’. Consecutively, the spot to surface Xtension was used to quantify the amount of kinesin motors on aforementioned surface areas.

### Total internal reflection (TIRF) microscopy & kinesin velocity analysis

GFP-tagged kinesin motors were recorded using a Zeiss Elyra PS1 (Zeiss) microscope with temperature control for activity recordings at 37°C. A 488 CW laser was used for excitation in combination with a Plan Apochromat 100x 1.46 NA Oil objective and CCD camera (Andor iXon DU-897 512×512). Kymographs were produced using the KymographBuilder plugin (Hadrien Mary, ImageJ) and stationary motors were removed by filtering in the frequency domain as shown in Supplementary Figure 6. Velocities were calculated using the Directionality plugin (ImageJ) written by Jean-Yves Tinevez^36^.

### Drosophila stocks and pharmacological treatment

All flies were kept on standard corn meal and sugar cane syrup at 25°C. Fly stocks used: w[1118]; P{y[+t7.7] w[+mC]=GMR57C10-GAL4}attP2 obtained from BDSC and y[*] w[*]; P{w[+mC]=UAS-tdTomato.mito}2 obtained from Kyoto stock center. For pharmacological treatment, flies of the appropriate genotype were collected within 8 hours from eclosion and kept on petri dishes with 20% sucrose and 1% agarose for 3 hours at 25°C. Starved flies were then moved to tubes with standard corn meal and sugar cane syrup supplemented with taxol (10 nM, 100 nM, 1 µM or 10 µM, Paclitaxel Cytoskeleton Inc.) or DMSO (0.01%, Cytoskeleton Inc.). Recordings of mitochondrial transport were performed after 24-28 or 48-52 hours of continuous drug exposure.

### Spinning disk *in vivo* mitochondrial transport recording & analysis

Flies were mounted in oil (refractive index 1.334, Zeiss Immersol) between glass coverslips spaced with double-sided tape as shown in Figure 6C and described previously ^37^. Mitochondrial transport recordings were performed on an inverted spinning disk microscope (Nikon Ti-Andor Revolution – Yogokawa CSU-X1 Spinning Disk) fitted with a Nikon 60x objective (Plan Apo, NA 1.27, W) and incubation chamber (Okolab, 25°C). TdTomato was excited with 561 nm laser light and a dual-band bandpass filter was used for emission (FF01-512/630-25, Laser 2000). The 10-minute long transport recordings consisted of stacks taken every 2 seconds and 10 micrometer thick to account for wing movement. Mitochondrial time lapse recordings were registered using the StackReg plugin (ImageJ) and in-house Igor pro code was used to generate kymographs and analyze mitochondrial transport time lapse recordings as described previously ^35,38,39^.

### Statistics

Graphpad Prism was used for statistical analysis: p < 0.05 (*), p < 0.01 (**), p < 0.001 (***) and bar graphs represent mean values with standard error of the mean. Shapiro-Wilk normality tests were used to assess the normal distribution of the data. T-tests were conducted as a two-sided test.

## Results

### Increased density of elongated GTP-tubulin dimers in axons is associated with slower anterograde transport

We first characterized the distribution of GTP- and GDP-bound tubulin dimer ratios in axonal and dendritic projections of hippocampal neurons. A general α-tubulin antibody was used to label all microtubules in combination with an antibody that recognizes specific tubulin dimers in a GTP-bound conformation. Dendritic and axonal processes were identified by immunohistochemistry (Fig. 1A). As previously shown ^21^, we confirmed that the relative GTP-bound tubulin content is higher in axons even though more microtubules are present in dendritic processes (Fig. 1B). To explore and study the effect of a higher GTP-tubulin density in axons on mitochondrial transport, we performed live mitochondrial transport imaging followed by post-hoc immunohistochemistry to distinguish the dendritic and axonal compartment as well as to determine the GTP-bound tubulin density (Fig. 1D, E). Dendritic processes have more motile mitochondria compared to axons, but the relative distribution of antero- and retrograde mitochondria is similar between both processes (Fig. 1F, G). To determine if the GTP- to GDP-tubulin ratio is related to mitochondrial transport velocity, we correlated GTP-tubulin density in all neuronal processes with transport velocities of the mitochondria present. We found a negative correlation for anterograde but not retrograde transport velocity, implying that GTP-tubulin dimers specifically slow down anterograde mitochondrial transport (Fig. 1H). A comparison of mitochondrial transport velocity in axonal and dendritic compartments revealed that anterograde velocity is significantly lower in axons, while retrograde transport is not different (Fig. 1I). These results suggest that the increased GTP-tubulin density in axonal projections slows down anterograde mitochondrial transport (Fig. 1J).

**Figure 1.**
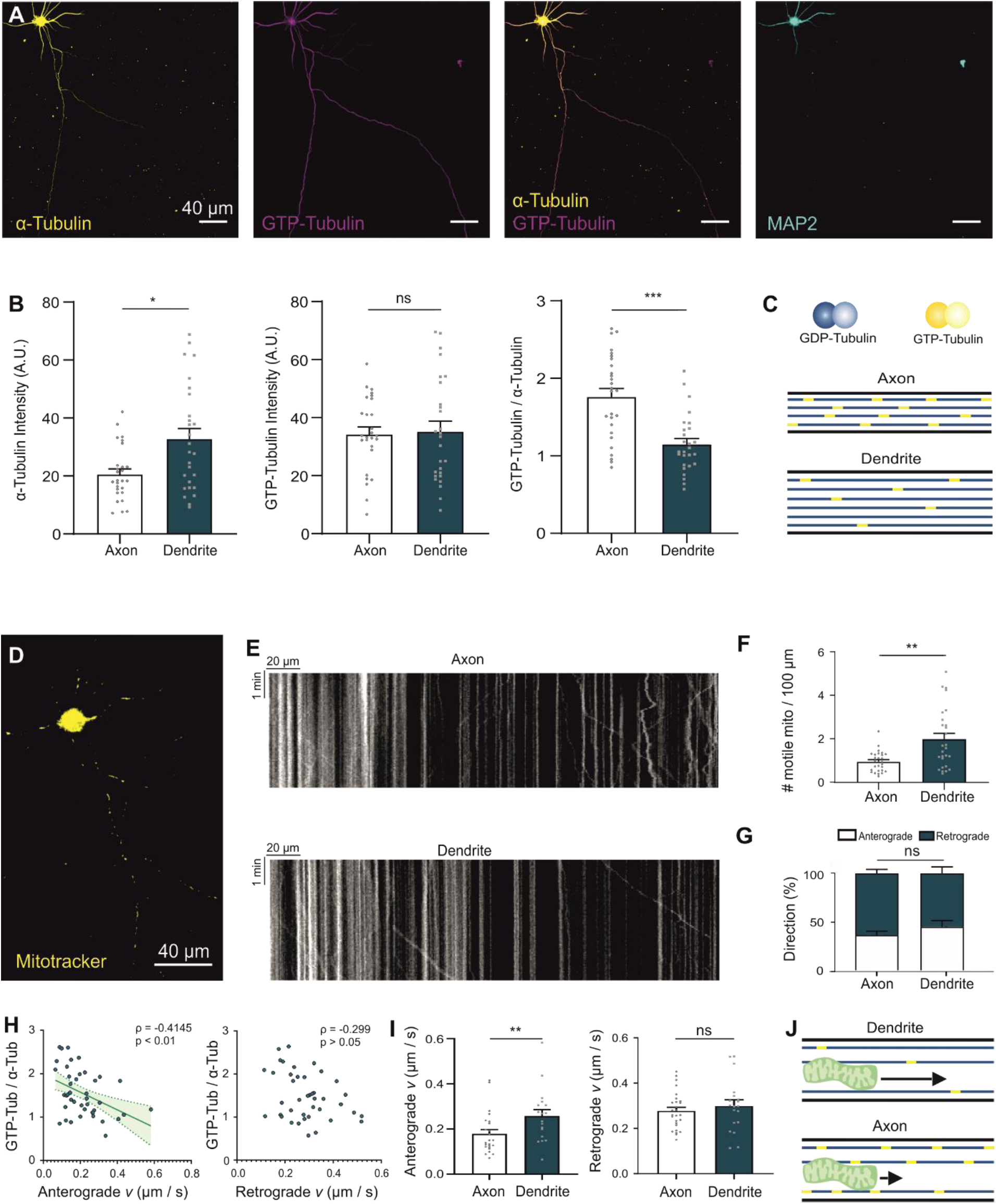
Increased density of GTP-tubulin dimers in axons negatively correlates with anterograde transport velocities. (A) Immunohistochemical staining of α-tubulin (yellow), GTP-bound tubulin dimers (magenta) and MAP2 (cyan) in hippocampal cultures at 7 DIV. (B) Quantification of the staining intensity of α-tubulin (n=27 cells from 3 independent experiments; * p<0.05 Mann Whitney test), GTP-bound tubulin (n=27 cells from 3 independent experiments; Unpaired two-tailed t-test) and their ratio (n=27 cells from 3 independent experiments; *** p<0.001 Unpaired two-tailed t-test) in either axonal or dendritic processes. (C) Schematic representation of the presence of GTP-bound tubulin islands in dendrites and axons. (D) Hippocampal neuron loaded with mitotracker red to visualize mitochondria. (E) Kymographs of axonal and dendritic projections. (F) Quantification of the number of motile mitochondria per 100 micrometer (n=27 cells from 3 independent experiments; ** p<0.01 Mann Whitney test). (G) No differences in directionality were found between axonal and dendritic motile mitochondria (n=27 cells from 3 independent experiments; Mann Whitney t-test). (H) A negative correlation was found between anterograde transport velocities and the amount of tubulin dimers in a GTP-bound conformation (n=42 neuronal processes from 3 independent experiments; ** p<0.01 Spearman correlation (- 0.4145), scatterplot ± 95% confidence interval) but not retrograde velocities and GTP-tubulin density (n=41 neuronal processes from 3 independent experiments; Spearman correlation (- 0.299)), derived from all neuronal projections. (I) Anterograde transport velocities were significantly higher in dendritic processes (n=24 cells from 3 independent experiments; ** p<0.01 Mann Whitney test) while retrograde transport velocities were not significantly different between dendrites and axons (n=27 cells from 3 independent experiments; Unpaired two-tailed t-test). (J) Schematic representation of suggested hypothesis that GTP-tubulin islands along microtubules could sterically hinder mitochondrial transport by increased straightness of the microtubule network.

### Taxol decreases anterograde mitochondrial motility but increases its velocity

To further understand if molecular changes in the tubulin dimer could alter mitochondria transport, taxol was used in neuronal cultures to increase the amount of elongated tubulin dimers. Treatment with taxol also reduces the number of motile mitochondria (Fig. 2A, B). Interestingly, changing the molecular conformation of the tubulin dimer has a specific effect on anterograde transport as taxol only decreased mitochondrial motility in the anterograde direction (Fig. 2C). Furthermore, while the amount of tubulin dimers in a GTP-bound conformation negatively correlates with average anterograde transport velocity in control cells (Fig. 1H), increasing the amount of elongated tubulin dimers beyond physiological levels with taxol leads to an increase in anterograde but not retrograde transport velocities (Fig. 2D). To exclude the possibility that the taxol effect was indirect, and a consequence of mitochondrial dysfunction, we performed an electron flow assay (Supplementary figure 1) on isolated mitochondria of mouse brain tissue incubated with taxol or DMSO. The functionality was similar and no significant changes in respiratory complex activity were detected between both groups. As the main difference in antero- and retrograde transport lies within the specificity of the motor proteins they employ – kinesin and dynein respectively –, we furthermore ensured that taxol had no effect on the binding of the kinesin motor machinery, as this would halt anterograde transport. To this end, we co-expressed two optogenetic constructs (TOM20-mCherry-LOVpep and Kif5b-GFP-ePDZb1), which allowed us to link mitochondria directly to kinesin motors upon illumination with blue light without the need to bind regulatory proteins (Supplementary figure 2). Upon stimulation of transport, taxol-treated cells still show decreased anterograde motility, indicating that taxol’s effect on transport does not act via the Miro-Milton regulator complex.

**Figure 2.**
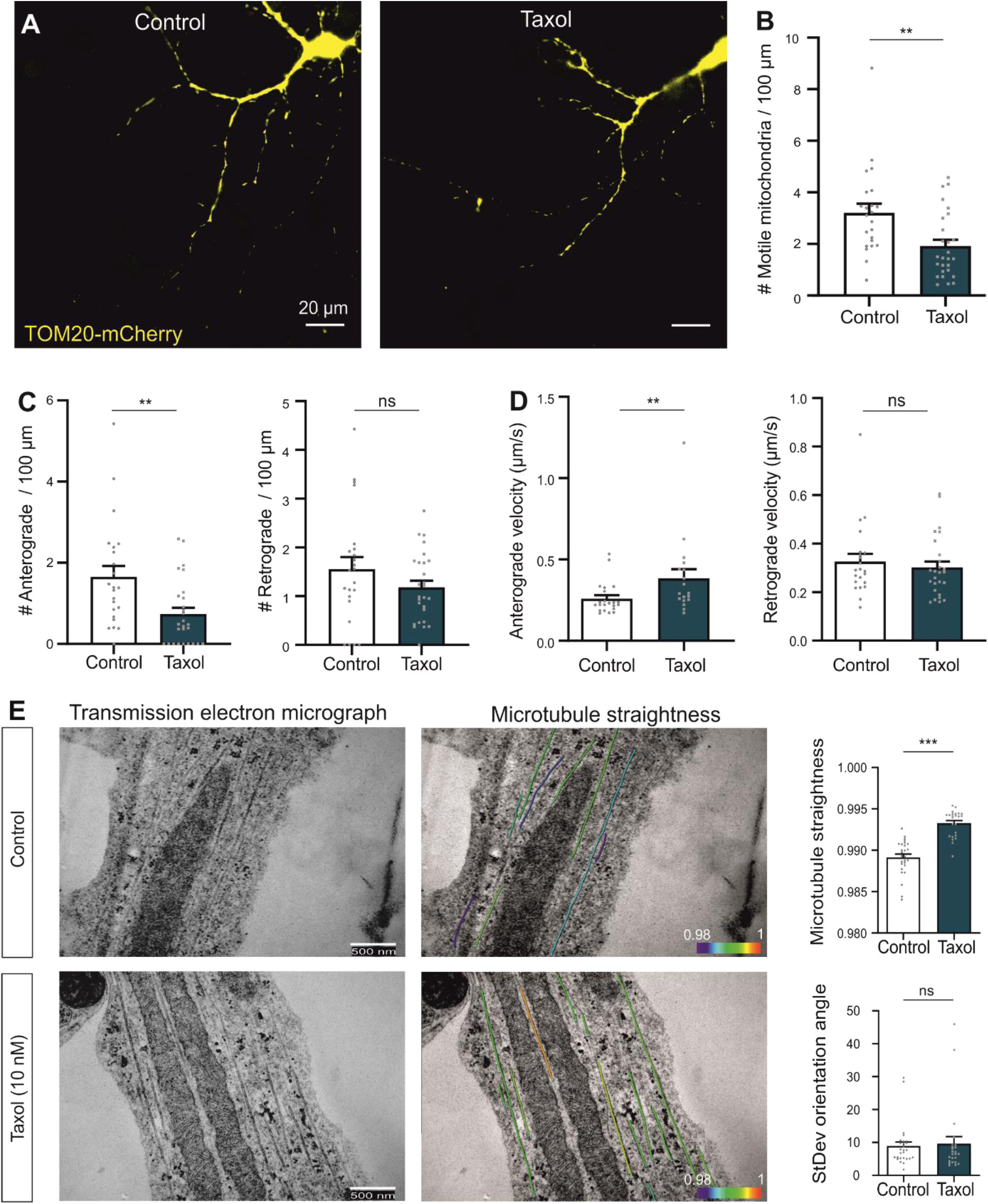
Taxol decreases anterograde mitochondrial motility but increases anterograde velocity and microtubule straightness. (A) Hippocampal neurons transfected with TOM20-mCherry, incubated with DMSO (Control) or taxol (4h, 10 nM). (B) Quantification of the number of motile mitochondria per 100 micrometer shows a significant reduction in motile mitochondria in taxol-treated cells compared to control neurons (n=27 neuronal processes from 3 independent experiments; ** p<0.01 Mann Whitney test). (C) The reduction in motile mitochondria numbers upon addition of taxol is due to a significant loss of anterograde moving mitochondria (n=27 neuronal processes from 3 independent experiments; ** p<0.01 Mann Whitney test) but not a loss of retrograde moving mitochondria (n=27 neuronal processes from 3 independent experiments; Unpaired Welch two-tailed t-test). (D) Although fewer motile mitochondria were present, a significant increase in anterograde but not retrograde velocities in taxol-treated cells was observed (n=22 neuronal processes from 3 independent experiments; ** p<0.01 Mann Whitney test). (E) Transmission electron micrographs of neurons treated with DMSO (Control) or taxol (10 nM, 4 h) with overlays of manually specified microtubule structures. Microtubule straightness was significantly increased (n=24 neuronal processes from 3 independent experiments; *** p<0.001 Mann Whitney test) in taxol-treated cells while the standard deviation of the orientation angles was not significantly different (n=24 neuronal processes from 3 independent experiments; *** p<0.001 Mann Whitney test).

### Elongated tubulin dimers increase microtubule straightness

While the increased density of GTP-tubulin dimers in axonal microtubules is beneficial for strengthening axon structure, we hypothesized that their presence could also render microtubules less flexible. As a consequence, axonal mitochondria would experience more resistance while being transported within the microtubule bundle, which, if true, could explain the decreased anterograde velocities (Fig. 1J). To determine the effect of the GTP-tubulin conformation on microtubule straightness in neuronal processes, we used taxol (10 nM) locking the tubulin dimer in a similar elongated conformational state as GTP-tubulin. Using transmission electron microscopy of longitudinally cut axons, we found that taxol-treated neurons contained microtubules that were more straight as compared to control samples while the standard deviation of the orientation angle was not significantly different, indicating that the microtubules were aligned along the length of the axons in the same way as in control neurons (Fig. 2E).

### Nano-positioning of mitochondria at the rim and within microtubule bundles

Although microtubule straightness is increased upon incorporation of elongated tubulin dimers, their rigidity would only hinder mitochondrial transport if mitochondria are also transported within the microtubule bundle rather than at the rim. As the exact nano-positioning of mitochondria during axonal transport is not known, we set out to study mitochondrial positioning during transport. To this end, hippocampal neurons were loaded with Mitotracker and SiR Tubulin to visualize mitochondrial transport and microtubules respectively using live STED microscopy (Supplementary Figure 3), revealing a subset of motile mitochondria indeed transported within the microtubule bundle (Fig. 3A). To further quantify mitochondrial localization, we added paraformaldehyde to the cells during mitochondrial transport recordings, concurrent with detection of mitochondrial mobility. This allowed identifying motile and stationary mitochondria during post-hoc immunohistochemistry (Fig. 3, Supplementary Figure 4A-G). We proceeded with 3D-STED microscopy on the fixed samples which enabled full 3D characterization of the mitochondrial localization relative to the microtubule network. The percentage of the optical surface overlap of microtubules per mitochondrion, hereinafter referred to as a surface^MITO-MT^ overlap, was calculated as a measure for mitochondrial confinement within the microtubule bundle. We found a median surface^MITO-MT^ overlap of 52.4%, indicating that a significant proportion of mitochondria is transported within the microtubule bundle (Fig. 3H).

**Figure 3.**
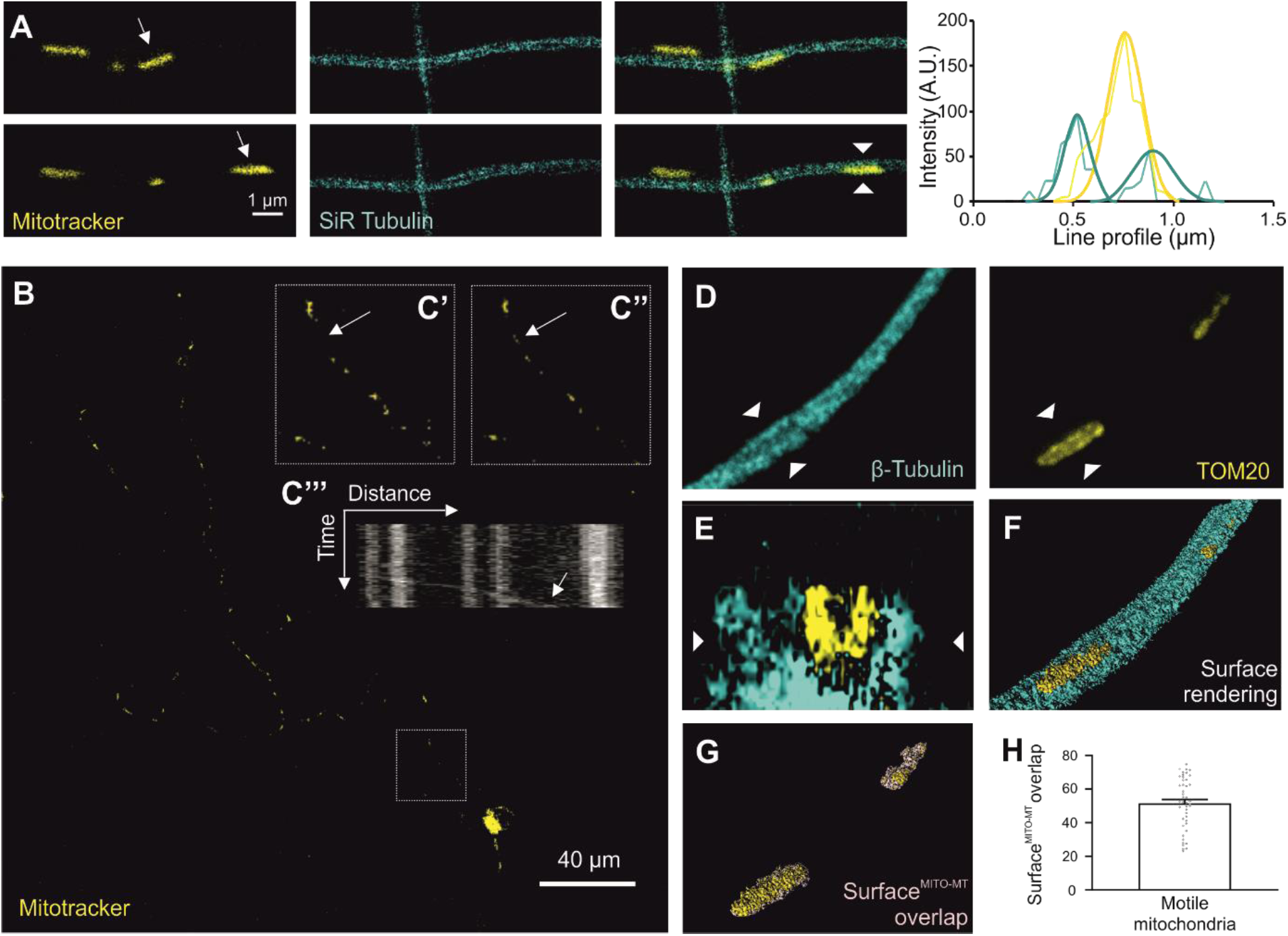
Nano-positioning of mitochondria at the rim and within microtubule bundles. (A) Live two-color STED time lapse recordings of hippocampal neurons loaded with mitotracker green (mitochondria, yellow) and SiR Tubulin (microtubules, cyan). A line profile perpendicular to the direction of transport (arrowheads) of a motile mitochondrion (arrows) is shown with the corresponding intensity profile for the tubulin (cyan plot and overlaid Gaussians) and mitochondrion (yellow and overlaid Gaussian) signal. (B) Mitotracker red was loaded onto hippocampal neurons before mitochondrial transport recordings. Paraformaldehyde was added during recordings to halt mitochondrial transport and all other intracellular processes. Kymograph indicating an anterograde motile mitochondrion as seen moving in panels (C’ and C’’, arrows). (D) After immunohistochemistry (TOM20 and ß-tubulin), z-stacks were recorded with a STED microscope. (E) Cross section of the z-stack. (F) Surface rendering of mitochondria and tubulin was performed followed by (G) calculation of the surface^MITO-MT^ overlap. (H) Quantification of the interface percentage of motile mitochondria with tubulin (n=44 mitochondria from 3 independent experiments).

**Figure 4.**
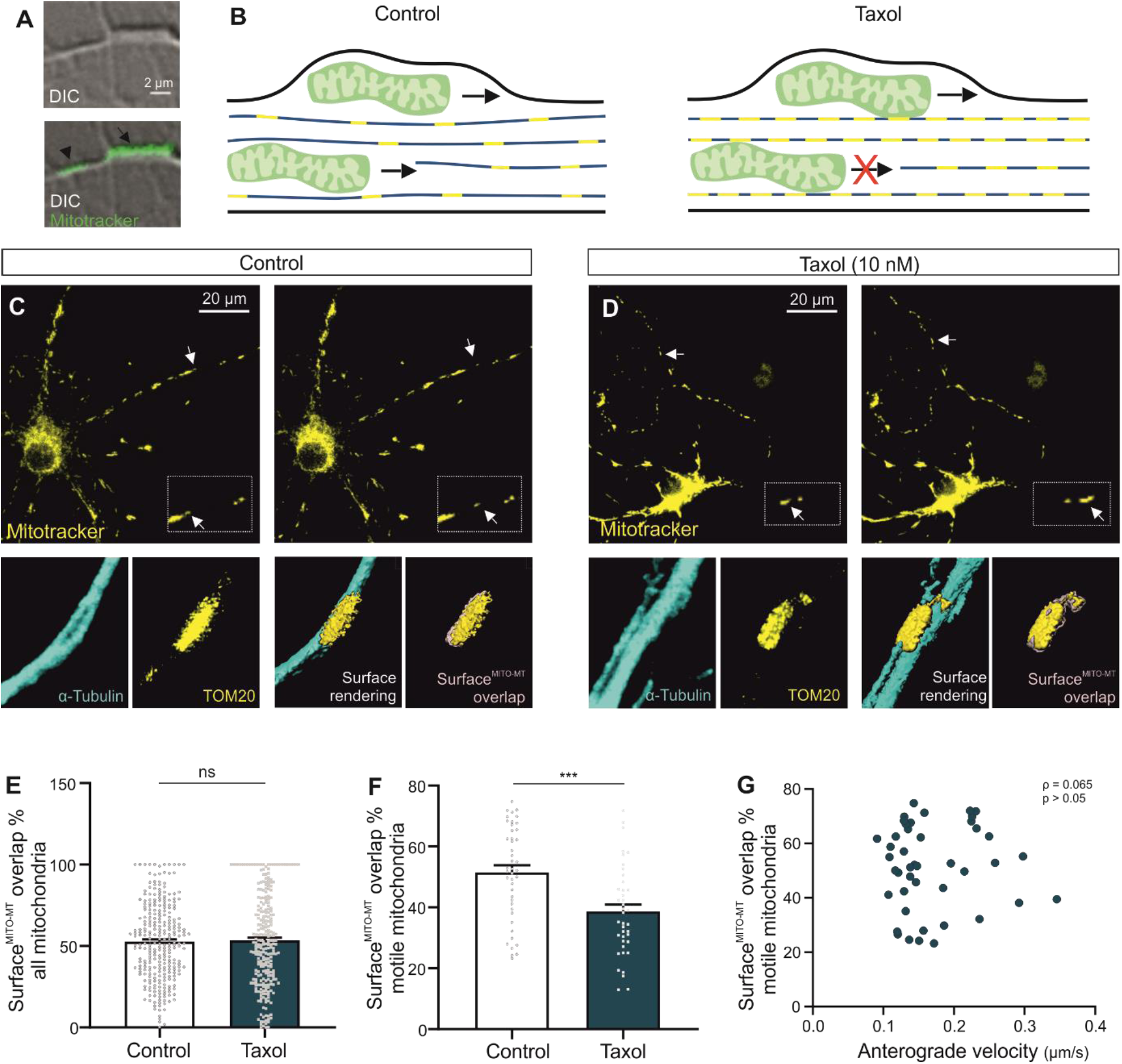
Excess of elongated tubulin dimers hinders mitochondrial transport within the microtubule bundle. (A) Differential interference contrast (DIC) image of a mitochondrion travelling between the microtubule bundle (arrowhead) or between the microtubule bundle rim and plasma membrane (arrow). (B) Schematic representation indicating that mitochondria between the microtubule bundles are no longer motile with increasing density of elongated tubulin while mitochondria at the rim of the microtubule bundle are still transported. (C) Hippocampal neurons were loaded with mitotracker after incubation with DMSO as control or (D) taxol (10nM, 4h). During transport recordings, paraformaldehyde was added to the imaging buffer at the moment an anterograde motile mitochondrion was observed (arrows). After fixation and immunohistochemistry using antibodies for β-tubulin and TOM20, z-stacks were recorded using STED microscopy. Surface rendering and calculation of the surface^MITO-MT^ overlap percentage. (E) Quantification of the surface^MITO-MT^ overlap of all mitochondria does not reveal a difference between control and taxol-treated samples (n=264 mitochondria from 3 independent experiments; Mann Whitney test). (F) A significant reduction in surface^MITO-MT^ overlap of anterograde motile mitochondria is measured when comparing control and taxol-treated cells (n=39 mitochondria from 3 independent experiments; *** p<0.001 Mann Whitney test). (G) No significant correlation was found between anterograde transport velocities and surface^MITO-MT^ overlap in control cells (n=44 mitochondria from 3 independent experiments; Spearman correlation (0.065)).

### Excess of elongated tubulin dimers hinders mitochondrial transport within the microtubule bundle

We hypothesized that the loss of motile mitochondria is mainly due to steric hindrance owing to the increased microtubule straightness induced by taxol (Fig. 4A, B). Therefore, only mitochondria transported between the microtubules would become immotile while mitochondria at the rim are still transported after taxol treatment. To test this, mitotracker was added to hippocampal cultures after incubation with taxol or DMSO (Fig. 4C, D). Cells were fixed on stage concurrent with detection of mitochondrial motility in the anterograde direction as shown previously (Supplementary Figure 4). Using STED microscopy on the fixed samples, no significant difference was detected in surface^MITO-MT^ overlap of all mitochondria (both motile and stationary) between control and taxol-treated samples (Fig. 4E). However, when only the motile mitochondrial fraction was considered, we observed a significant decrease in surface^MITO-MT^ overlap in taxol-treated cells (Fig. 4F). The increased microtubule straightness in taxol-treated cells thus halts specifically those motile mitochondria that are transported within the microtubule bundle as illustrated in Figure 4B. As the (remaining) motile fraction of mitochondria in taxol treated cells appears to represent the mitochondrial fraction that travels along the rim of the microtubule bundle, where less steric hindrance is expected, their increased anterograde velocities could be related to their specific position. However, there was no significant correlation between the surface^MITO-MT^ overlap and anterograde velocities in control cells (Fig. 4G). These results indicate that while the increased straightness of the elongated tubulin conformation specifically halts motile mitochondria transported within the microtubules bundle, the velocity of the remaining motile mitochondria is not determined by their positioning with respect to the microtubule network.

### Kif5B preferentially links mitochondria to elongated tubulin and increases mitochondrial velocity

As the increased anterograde mitochondrial velocity in the presence of taxol cannot be explained simply by their localization with respect to the microtubule network (Fig. 4G), we set out to determine whether the effect is linked to the kinesin motor involved in anterograde mitochondrial transport: KIF5B. Multicolor STED microscopy was used to obtain sufficient resolution to detect KIF5b motor proteins on the motile mitochondrion’s surface (Fig. 5A, Supplementary Figure 5). Surfaces of motile mitochondria, the surrounding microtubule network and their surface^MITO-MT^ overlap were rendered while KIF5B motors were rendered using the spot detection feature of Imaris (Fig. 5B, C). The amount of KIF5B motors on the total mitochondrial surface was not significantly different between control and taxol-treated cells (Fig. 5D). Interestingly, the amount of KIF5B motors present on the surface^MITO-MT^ overlap, was significantly increased in taxol-treated neurons (Fig. 5E). The increase in bound motor proteins could increase anterograde transport velocities owing to increased pulling force. However, we found no significant correlation between anterograde velocities and the amount of kinesin motors at the surface^MITO-MT^ overlap (Fig. 5F), arguing that the increase in kinesin motors linked to mitochondria does not influence mitochondrial transport velocities. Besides the number of bound motors, also the processive motion of the motor protein could influence transport velocity. We therefore investigated KIF5B processive motion on distinct molecular tubulin conformations in neurons expressing Kif5b-GFP, partially bound to mitochondria (TOM20-mCherry), using total internal reflection (TIRF) microscopy (Fig. 5G, H and Supplementary Figure 6) and found that kinesin motors move significantly faster in taxol-treated cells (Figure 5I). These results indicate that the remaining motile mitochondrial fraction is located at the microtubule rim, is transported faster in the anterograde direction not because of the increased number of bound kinesin motors but owing to increased processive motion of the kinesin motor itself (Figure 5J).

**Figure 5.**
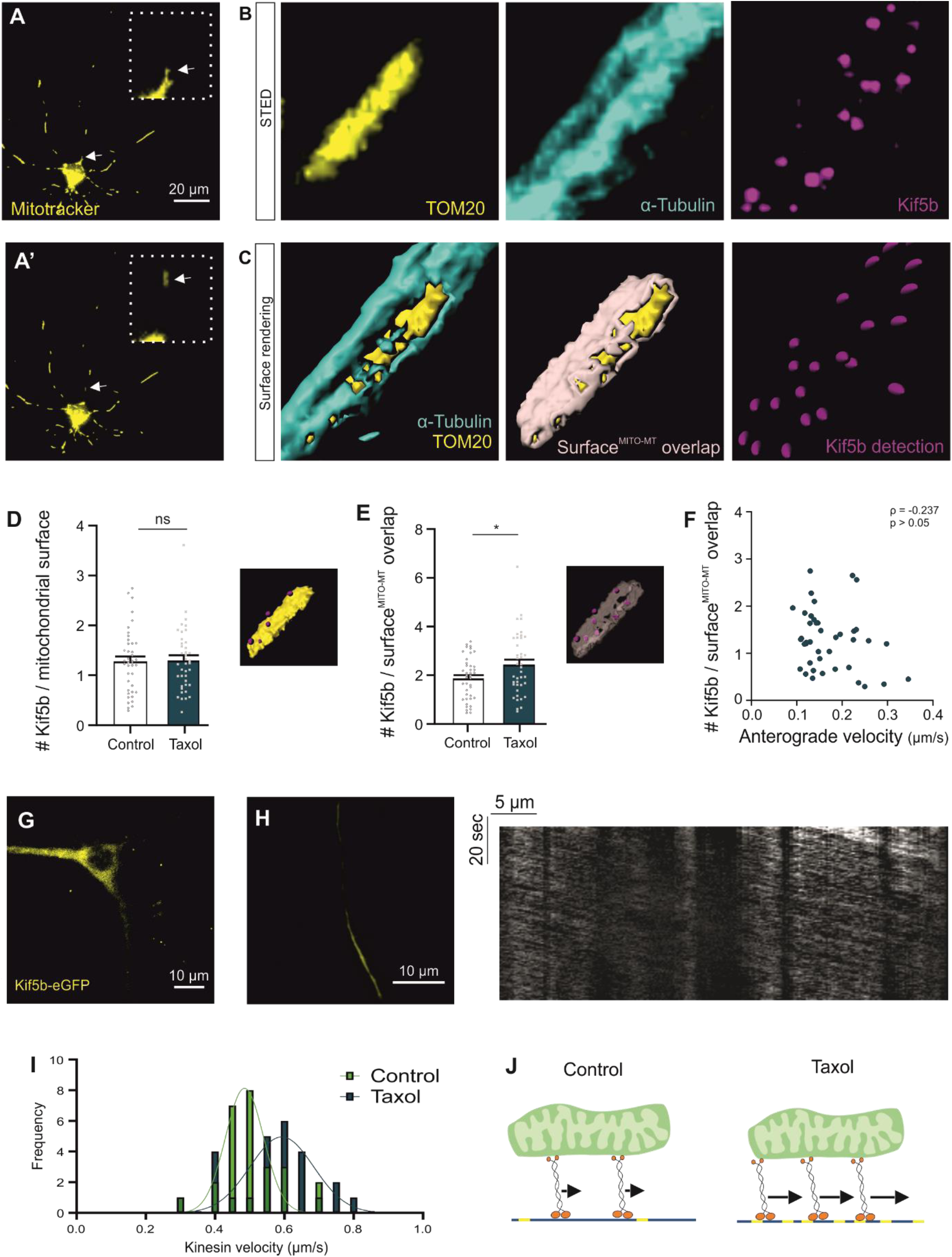
Kif5B preferentially links mitochondria to elongated tubulin and increases mitochondrial velocity. (A) Hippocampal neurons loaded with mitotracker after incubation with DMSO as control or taxol (10nM, 4h). During transport recordings, paraformaldehyde was added the moment an anterograde motile mitochondrion was observed (arrows). (B) 2D STED microscopy was used to record Z-stacks on the fixed samples using antibodies for α-tubulin, TOM20 and KIF5B. (C) Surface rendering and calculation of the surface^MITO-MT^ overlap. Kinesin motors were rendered using the spot detection module (magenta). (D) Quantification and comparison of the number of KIF5B spots on the entire mitochondrial surface shows no significant difference in control or taxol-treated cells (n=40 mitochondria from 3 independent experiments; Mann Whitney test). (E) Quantification of KIF5B spots on the surface^MITO-MT^ overlap shows significant increase in taxol-treated samples (n=40 mitochondria from 3 independent experiments; * p<0.05 Unpaired Welch two-tailed t-test). (F) No significant correlation was found between anterograde transport velocities and the amount of KIF5B motor proteins on the surface^MITO-MT^ overlap (n=40 mitochondria from 3 independent experiments; Pearson correlation (−0.237)). (G) Kif5b-eGFP signal of hippocampal neurons transfected with TOM20-mCherry and Kif5b-eGFP constructs. (H) TIRF microscopy of kinesin motors using the Kif5b-eGFP construct and example kymograph to indicate kinesin motility. (I) Frequency distribution of KIF5B velocities in DMSO (control) or taxol-treated samples (10 nM, 4h) shows increased velocities in taxol-treated cells (n=27 neuronal fibers from 3 independent experiments; ** p<0.01 Gaussian Least squares fit). (J) Schematic representation that the amount of kinesin motors bound to both mitochondria and tubulin is increased in taxol-treated samples and their individual velocities (arrow) are increased.

**Figure 6.**
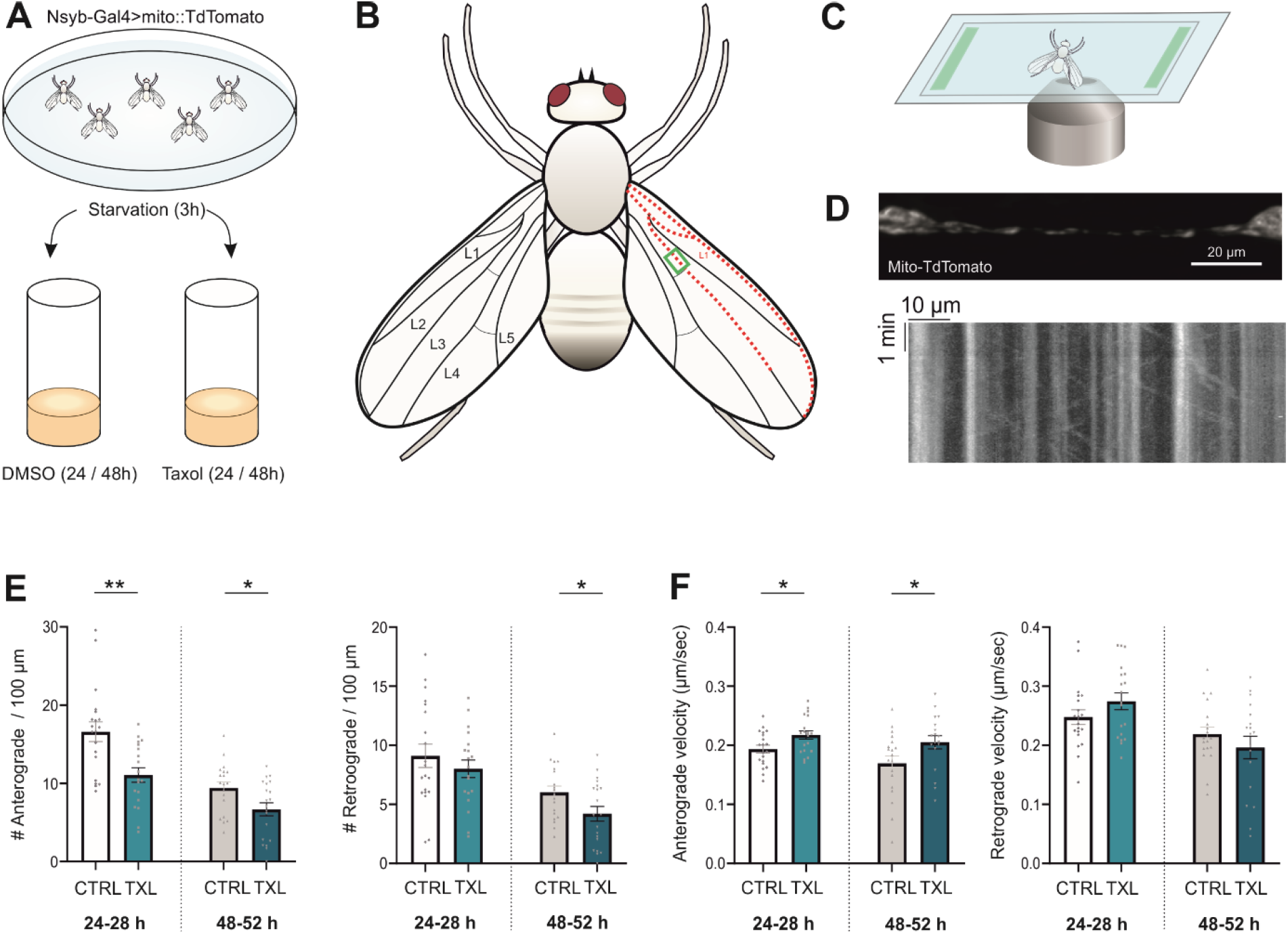
Taxol halts the majority of mitochondrial transport in the anterograde direction while increasing anterograde transport velocities in *vivo*. (A) *UAS-mito::tdTomato*/+; *Nsyb-Gal4/+* virgins were selected for pharmacological treatment within 8 hours from eclosion. After a 3 hour starvation period, flies were transferred to food vials containing regular food supplemented with DMSO as control or taxol (100 nM) for 24 or 48 hours before imaging. (B) Axonal projections and neuronal cell bodies were visible along the L1 and L2 veins in the wing, the branching region in L2 was used for transport recordings to reduce the number of axons and facilitate analysis. (C) Flies were mounted between two coverslips using double-sided tape and mitochondrial transport recorded using an inverted spinning disk microscope. (D) Sum projection of a mitochondrial time lapse recording and corresponding kymograph. (E) Quantification of the number of motile anterograde mitochondria per 100 micrometer shows a significant reduction in motile mitochondria in taxol (TXL)-treated flies compared to control neurons after 24h (n=19 flies from 3 independent experiments; ** p<0.01 Unpaired two-tailed t-test) and 48h (n=19 flies from 3 independent experiments; * p<0.05 Unpaired two-tailed t-test). The number of retrograde motile mitochondria remained unchanged after 24 h (n=19 flies from 3 independent experiments; * p<0.05 Unpaired two-tailed t-test) but significantly decreased after 48 hours taxol (n=19 flies from 3 independent experiments; * p<0.05 Unpaired two-tailed t-test). (F) Anterograde transport velocities were significantly increased after 24 h and 48 h taxol treatment while retrograde velocities remained unchanged compared to controls (n=19 flies from 3 independent experiments; * p<0.05 Unpaired two-tailed t-test, for each condition)

### Taxol halts the majority of mitochondrial transport in the anterograde direction while increasing anterograde transport velocities *in vivo*

Finally, we tested whether the conformation of tubulin dimers also regulates mitochondrial transport *in vivo. Drosophila m.* with neuronal TdTomato-tagged mitochondria were selected within 8 hours from eclosion and put on starvation for 3 hours followed by feeding on regular food supplemented with DMSO or 100 nM taxol (Figure 6A). After 24 and 48 hours of continuous taxol administration, flies were mounted in glass chambers and mitochondrial transport was recorded in the neuronal projections adjacent to vein L2 (Figure 6B-D). The taxol concentration was chosen based on a pilot experiment (Supplementary Figure 5) that assessed taxol intake and relevant changes in mitochondrial transport parameters using increasing taxol concentrations. We found a significant decrease in anterograde mitochondrial motility after 24 and 48 hours of continuous taxol administration, while retrograde transport was only significantly reduced after 48 hours (Figure 6E). Anterograde transport velocities were significantly increased after 24 and 48 hours taxol while retrograde transport velocities remained unchanged (Figure 6F). Thus, changing the tubulin dimer conformation regulates anterograde mitochondrial transport *in vivo* in a similar manner.

## Discussion

The central nervous system has a high-energy need and proper spatial and temporal distribution of the cell’s energy factories, mitochondria, are crucial ^8,40,41^. Failure to transport mitochondria has been linked to several neurodegenerative diseases including Parkinson’s disease, Alzheimer’s disease and Amyotrophic Lateral Sclerosis ^8,42,43^, which boosted studies on mitochondrial transport mechanisms. Apart from the identification of distinct motor protein families, also unraveling regulatory proteins such as the Miro-Milton complex and its calcium dependency was key for understanding transport in neurons ^44-47^. Changes at the level of the tubulin dimer can also influence intracellular transport as the nucleotide (GTP/GDP) bound to the tubulin dimer determines its conformation ^28,48^ and consequently affects motor protein activity ^21,25,26,49^. Drugs interfering with tubulin conformations such as taxol, a drug that favors GTP-tubulin and locks it in its elongated conformational state ^15,28,48,50^, have also been shown to disrupt transport dynamics ^30-32^. However, exactly how transport is affected upon changes in the conformational state of the tubulin dimer is not fully understood. Here, we describe how the tubulin dimer conformation regulates mitochondrial transport dynamics and show that this regulatory mechanism is susceptible to mitochondrial positioning along the microtubules.

In agreement with previous research, we first show that axonal projections have a higher GTP-tubulin density as compared to dendrites ^21^. Furthermore, transport dynamics differ in axons and dendrites with fewer motile mitochondria and reduced anterograde velocities in axons. A negative correlation between GTP-tubulin density and anterograde transport velocities in axons indicate a possible relationship between tubulin conformations and transport dynamics. To understand the regulatory effect of conformational changes in the tubulin dimer on mitochondrial transport, we compared mitochondrial transport parameters between control- and taxol-treated cells. While the number of anterograde motile mitochondria was significantly reduced upon taxol treatment, no change in the number of retrograde motile mitochondria was detected. This could be linked to the fact that kinesin motors bound to cargo have more difficulty traversing obstacles owing to their limited step-size and short neck-linker as compared to dynein motors ^51-54^. Increasing the amount of stable, elongated tubulin dimers in neurons also led to increased anterograde-but without affecting retrograde velocities. This is in line with the direction-dependent decrease in motile mitochondria and indicates that processivity of kinesin motors depends on the conformational changes in tubulin dimers.

The negative correlation between anterograde transport velocities and elongated tubulin density in neuronal projections in control cells and reduction of motile anterograde mitochondria in taxol-treated cells could be explained by steric hindrance. Indeed, we show that microtubules become more rigid upon incorporation of elongated tubulin dimers, as shown previously using *in vitro* polymerized microtubules ^48^. Furthermore, mitochondria are also being transported between these (rigid) microtubules and not solely at the rim (using live 2D and fixed 3D STED recordings. The presence of elongated rather than compacted tubulin dimers could thus control anterograde transport within the microtubule bundle.

We hypothesized that the opposite effects on mitochondrial transport upon increased elongated tubulin dimers, namely a halt in anterograde transport but an increase in anterograde velocities, could be linked to the mitochondrial localization within the microtubule network where its rigidity could influence a specific subpopulation of motile mitochondria. We show that the reduced motile fraction in taxol-treated samples is mainly located at the rim of the microtubule bundle, indicating that those mitochondria travelling between the rigid microtubule bundle become stationary. Interestingly, while the halting of mitochondria could be attributed to their location between rigid microtubules, the increased velocities observed at the rim could not be explained solely by mitochondrial positioning.

To elucidate why mitochondria at the rim of taxol-treated neurons move at increased velocities, we focused on kinesin motor proteins and their interaction with distinct molecular tubulin conformations. We found a significant increase in the amount of kinesin motors located on the surface overlap of specifically motile mitochondria and taxol-treated microtubules, in line with a previous study by Nakata *et al*. showing that kinesin-1 preferentially binds elongated GTP-bound tubulin dimers on *in vitro* polymerized microtubules ^21^. However, this increased amount of bound kinesin motors does not influence mitochondrial transport velocity as shown before using *in vitro* polymerized microtubules ^51,55^.

In a following set of experiments, we investigated whether the tubulin conformation influences kinesin processive motion by altering their velocity directly. We show that kinesin motors move faster on taxol-treated microtubules, explaining the increased mitochondrial anterograde velocity in taxol-treated cells which aligns with results reported using *in vitro* polymerized microtubules ^26^. We hypothesize that this faster stepping process can occur as the kinesin motor no longer has to pull the tubulin dimer in an elongated conformation ^25^. While most studies using *in vitro* polymerized microtubules agree with our findings in neuronal cultures, some studies concerning kinesin-1 binding affinity and velocity report no significant difference between microtubules consisting of the elongated – or compacted tubulin conformation ^56,57^. It should be noted that the latter studies compared similar conformational states of the tubulin dimer, i.e. microtubules polymerized with GMPCPP, a very slowly hydrolysable GTP analog, and taxol-stabilized microtubules. While these studies indicate the importance of stabilization methods for *in vitro* polymerized microtubule assays, they also provide further evidence of the conformational similarities between taxol-bound tubulin dimers and native GTP-bound tubulin as no kinesin-dependent differences between both microtubule populations were reported.

Finally, we confirmed that tubulin conformations are also relevant in regulating mitochondrial transport *in vivo*. To change the molecular conformation of tubulin dimers, we fed adult *Drosophila m.* taxol. Previous studies using oral administration of taxol in fruit flies are limited to larvae and use high concentrations that lead to neuronal degeneration ^58,59^. As we were mainly interested in early effects of taxol, preceding axonal degeneration, we first performed a pilot study using various low taxol concentrations. From the bell-shaped dose-velocity curve, we opted for 100 nM taxol for oral administration to ensure enough taxol is taken up while reducing possible adverse secondary effects linked to axonal degeneration. We found that taxol reduced anterograde motility both after 24 and 48 hours, while retrograde transport was decreased only after 48 hours, possibly a consequence of fewer mitochondria being present due to early onset reduced anterograde transport. We also note that mitochondrial transport in control flies decreased during maturation as shown previously ^60^. Furthermore, anterograde transport velocities are significantly increased after taxol administration for both timepoints while retrograde transport velocities remain unchanged.

In summary, we show that the molecular conformation of the tubulin dimer influences mitochondrial transport in two distinct ways depending on mitochondrial localization. On one hand, the increased straightness of microtubules enriched in elongated tubulin hinders specifically those mitochondria travelling within the microtubule bundle, possibly explaining why studies investigating CIPN linked to taxol treatment observe a decrease in motile organelles and vesicles ^30-32,61^. On the other hand, mitochondria mobile at the rim of the microtubule bundle are transported at increased velocities as the molecular conformation of tubulin affects kinesin binding and processive velocity in neuronal projections. As an increased density of elongated dimers leads to straighter microtubules, it is conceivable that processive motion of kinesin motors is mechanically more efficient resulting in increased transport speed. We conclude that the conformation of tubulin dimers within the microtubule lattice is an important regulator of intracellular transport, both *in vitro* and *in vivo*, and that the specific location of motile mitochondria is crucial for the regulatory effect. These findings are relevant not only to understand intracellular transport mechanisms but are also crucial for cancer research and more specifically how chemotherapeutic agents can interfere with neuronal processes such as intracellular transport.

## Acknowledgments

We would like to thank M. Moons for technical support and all current and past LENS members for scientific comments on the project. We thank Prof. J. Hendrix (and Dr. S. Duwé for assistance with TIRF recordings at the Dynamic Bioimaging Lab (BIOMED, UHasselt). We finally thank Prof. P. Agostinis for use of the Seahorse analyzer and Prof. L. Kapitein for the optogenetic constructs.

## Author contributions

Methodology: VVS, PVB. Investigation: VVS, FLC, YK, MD, TM. Data discussion: VVS, WB, PV, PVB. Writing – Original draft: VVS. Writing – Review & Editing: VVS, FLC, YK, MD, TM, PV, WB, PDK, PVB. Conceptualization: VVS, PVB. Funding: PV, PDK, PVB.

