## Supplemental figures for "Nano-positioning and tubuline conformation determine transport of mitochondria along microtubules"

For:

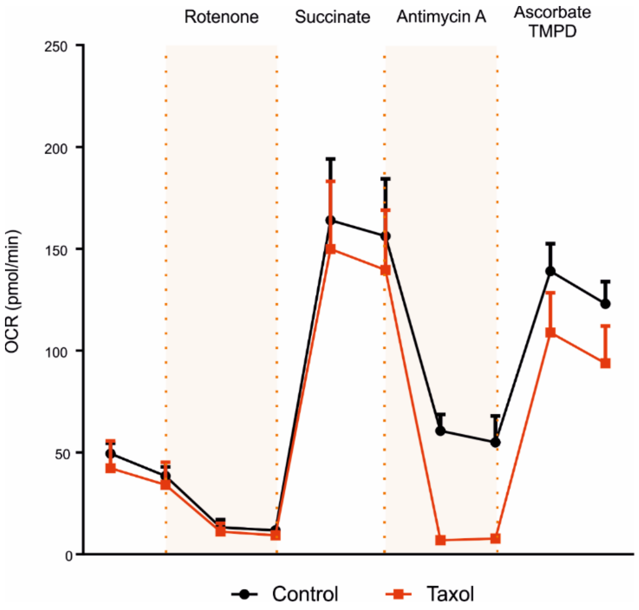


**Supplementary Figure 1. There is no change in oxygen consumption rate in isolated mitochondria incubated with taxol.**

Isolated mitochondria from hippocampal brain tissue were incubated in either DMSO (control) or taxol (10 nM, 1 h). For the electron flow assay, 20 µM rotenone, 100 mM succinate, 40 µM antimycin A and 100 mM ascorbate with 1mMB TMPD were used. No significant change in oxygen consumption rate (OCR) was detected (n=3 isolations; Two Way ANOVA with Bonferroni).

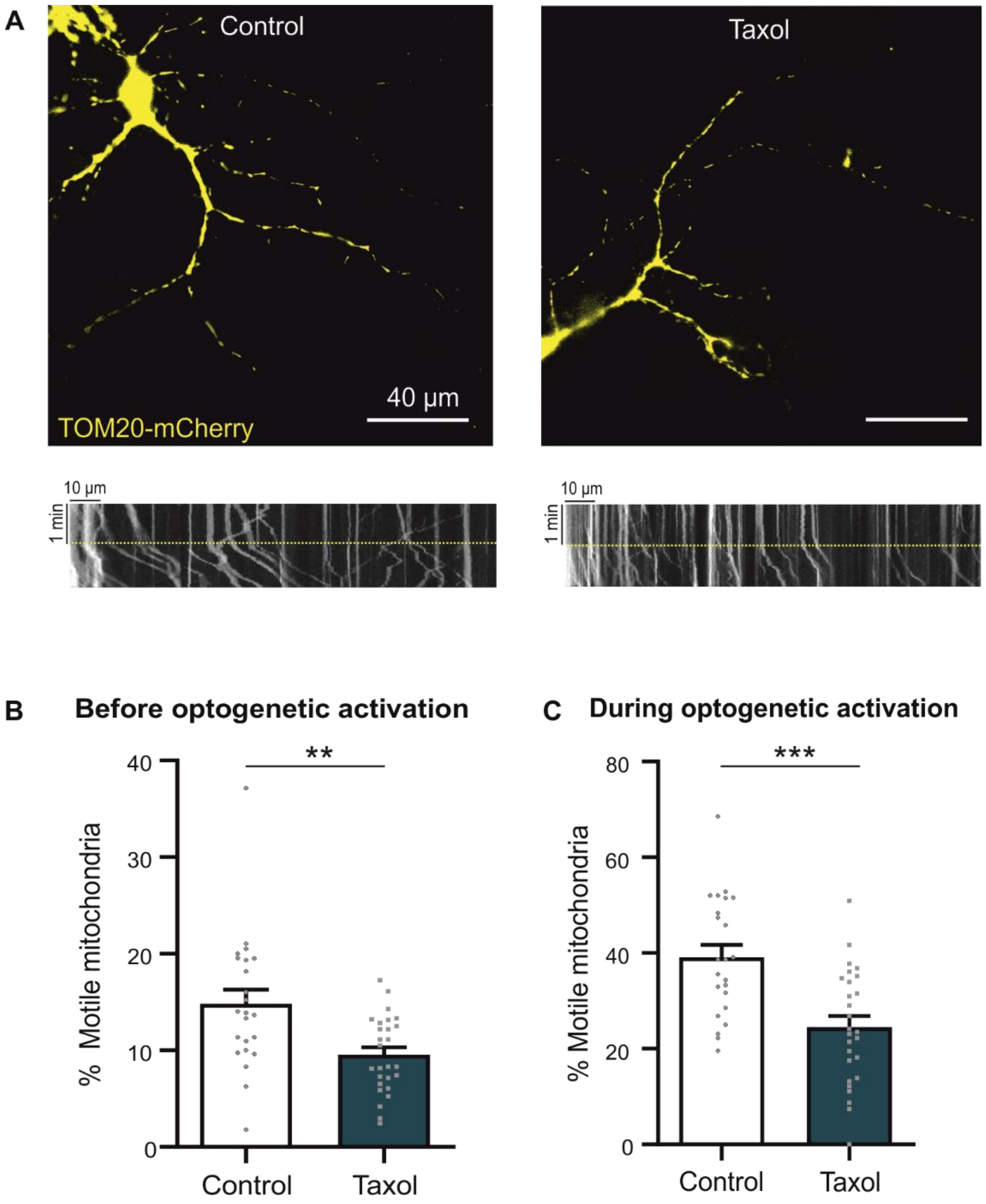


**Supplementary Figure 2. The reduction in motile mitochondria in taxol-treated cells is not linked to the kinesin-associated proteins required for transport.**

(A) Hippocampal neurons were transfected with two constructs: TOM20-mCherry-LOV and Kif5b-GFP-ePDZb1. Upon illumination with blue light (indicated by the yellow dotted line in the kymograph), the Light-Oxygen-Voltage (LOV)-domain and ePDZ1 domain heterodymerize, forcing a link between kinesin motors (KIF5B) and mitochondria (TOM20). (B) Before optogenetic activation, there is a significant reduction in the percentage of motile mitochondria in taxol-treated (10 nM, 4 h) cells compared to controls (n=27 axons from 3 independent experiments; ** $p<0.01$ Mann Whitney Test). (C) After optogenetic activation, the reduction in motile mitochondria after taxol incubation (10 nM, 4h) remains significant compared to control cells (n=27 axons from 3 independent experiments; *** $p<0.001$ Mann Whitney Test).

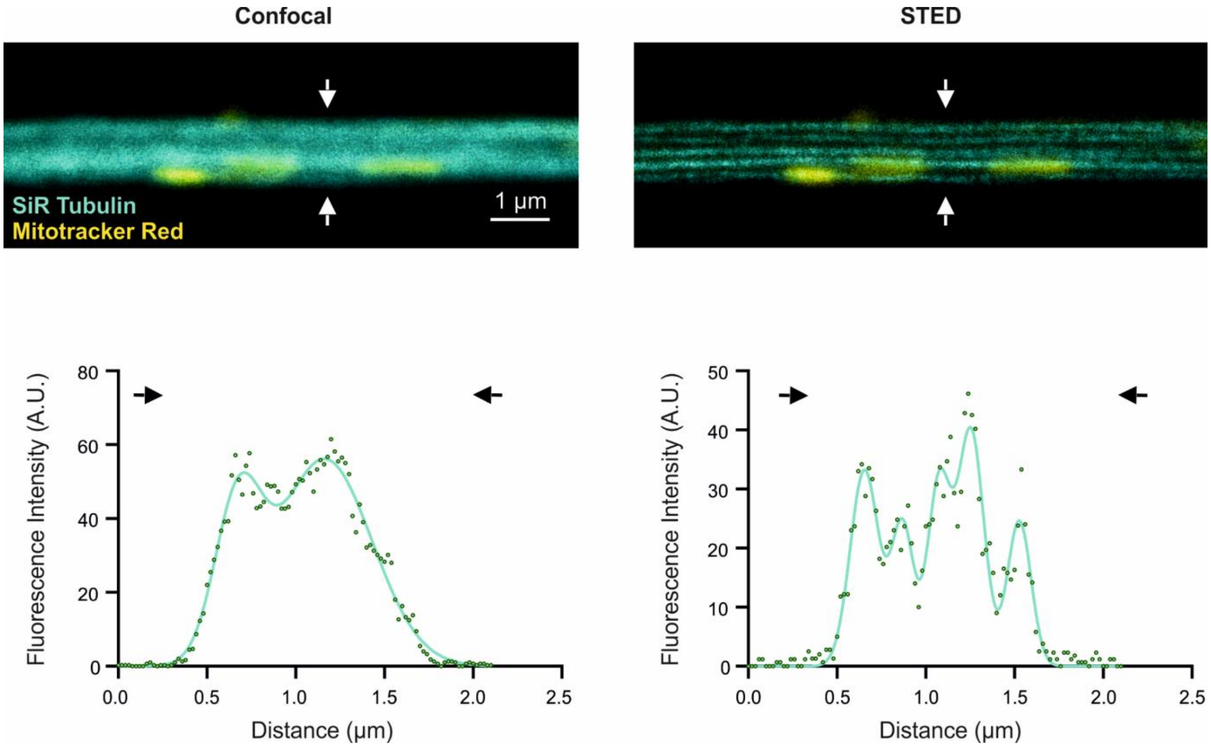


**Supplementary Figure 3. Super resolution live STED imaging of microtubules.**

Hippocampal neurons were loaded with mitotracker red (yellow) and SiR Tubulin (cyan). Mitochondria were recorded in confocal mode, microtubules in both confocal (left) and STED (right). Bottom graphs represent line profiles across the fiber (arrows) to indicate increased resolution (multiple Gaussian fit).

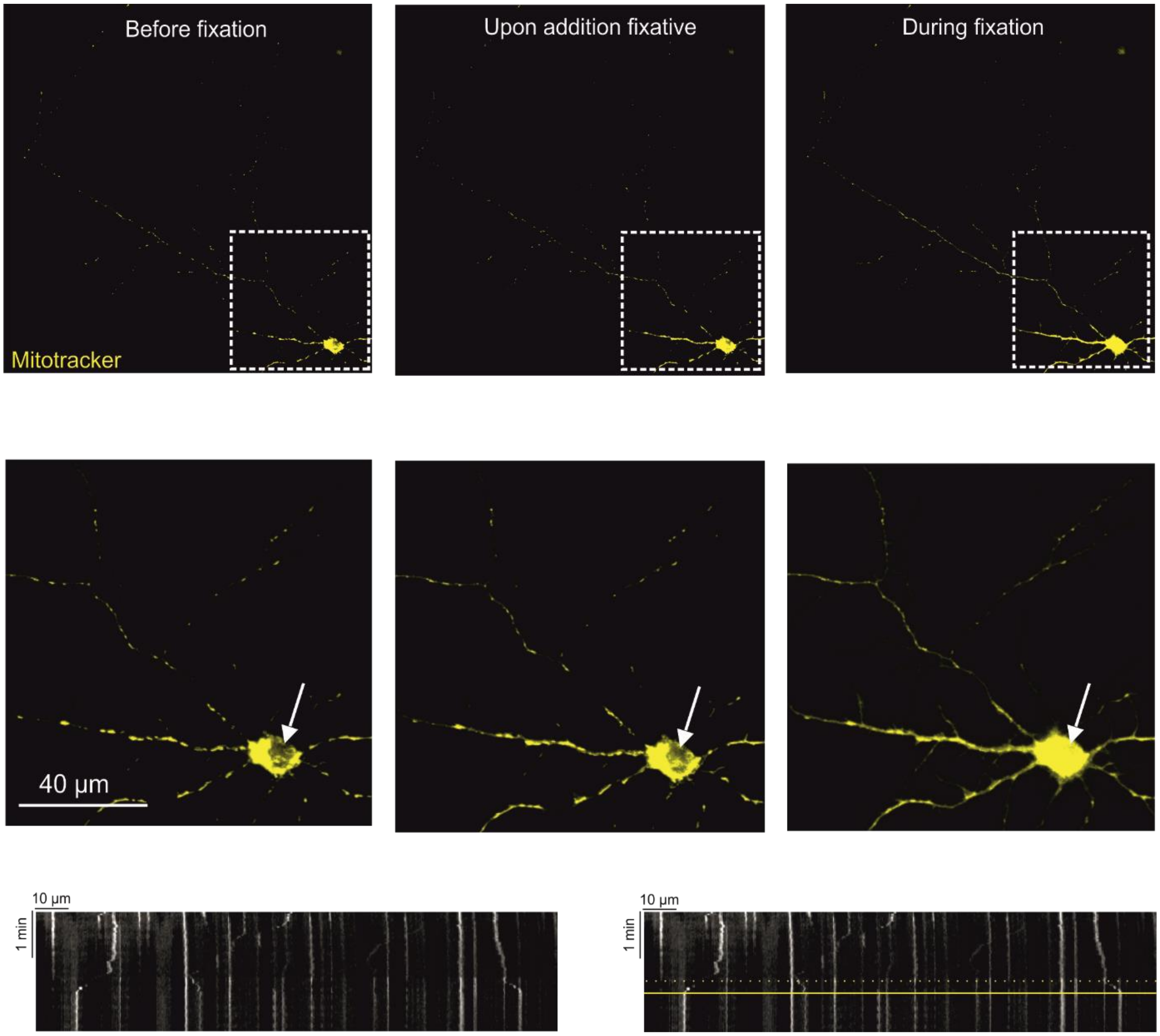


**Supplementary Figure 4. Addition of paraformaldehyde during transport recordings halts all mitochondria**

Mitochondrial transport recordings on mitotracker-loaded neurons before, upon and after addition of paraformaldehyde with resulting kymograph (bottom left). The effect of fixation is easily observed in the cell soma (arrow), where mitotracker is no longer accumulating in mitochondria but also in the nucleus. Upon addition of fixative, a small change in intensity is observed due to the altered buffer volume (dashed yellow line, bottom right kymograph) and all mitochondria halt as seen in the kymograph (yellow line, bottom right).

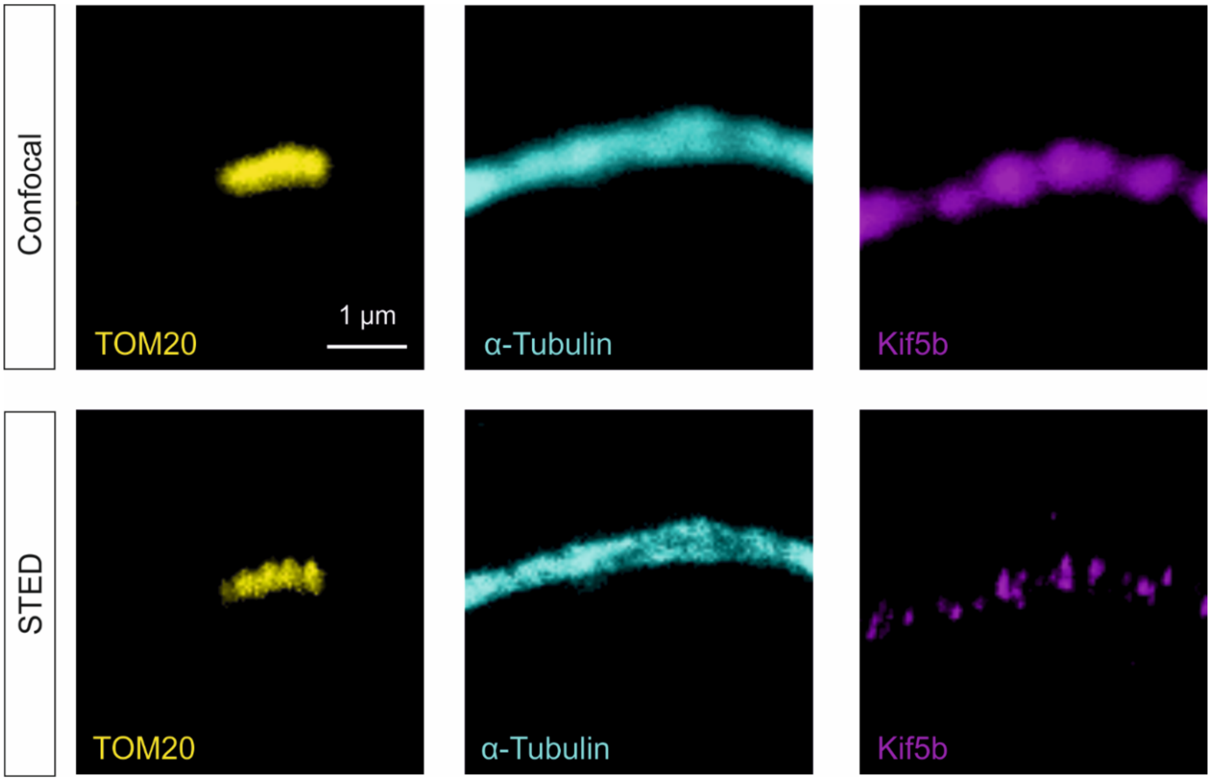


**Supplementary Figure 5. Increased resolution of STED microscopy allows visualization of kinesin motors.**

Confocal and STED recordings shown in the upper and lower panels respectively for TOM20 (yellow), α-tubulin (cyan) and Kif5b (magenta) immunostainings. Images represent maximum intensity projections of Z-stacks.

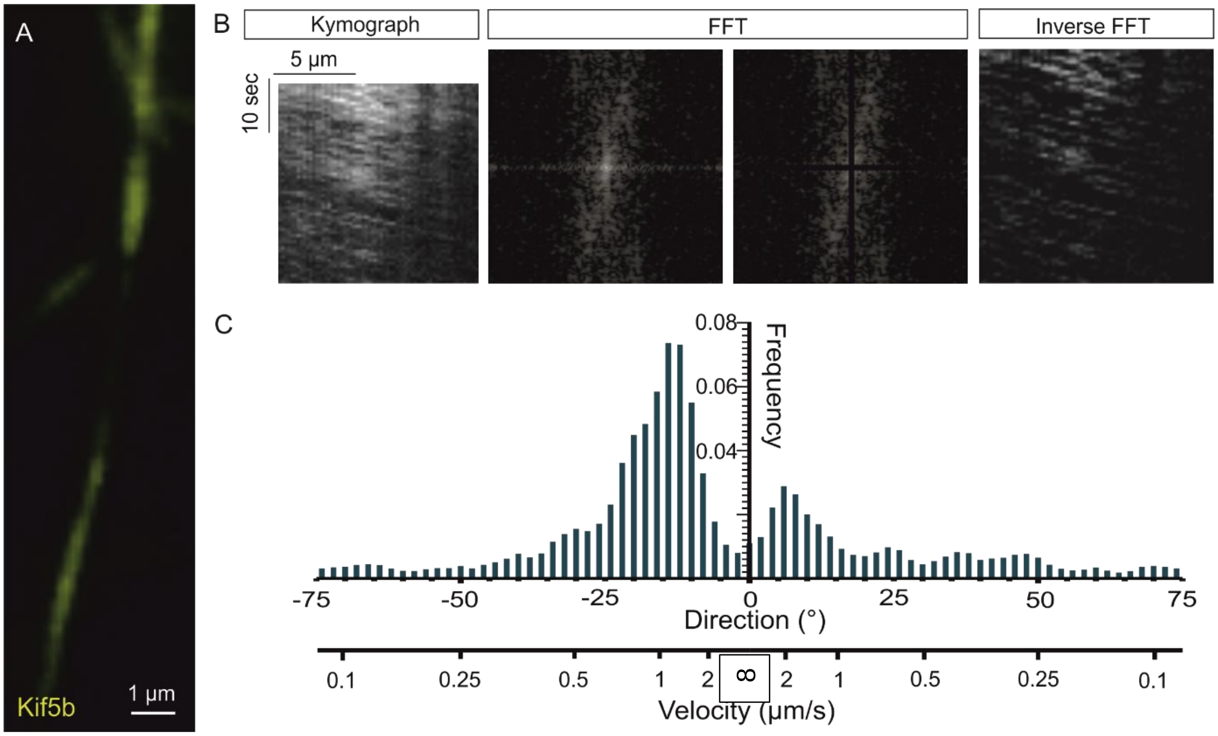


**Supplementary Figure 6. Post-processing of Kif5b kymographs and frequency distributions of directional angles related to kinesin velocities.**

(A) Maximum projection of TIRF time lapse recording in Kif5b-GFP-ePDZb1 transfected hippocampal neurons. (B) Kymograph and corresponding Fast Fourier Transform (FFT), after the removal of frequencies along the horizontal and vertical axis in the Fourier image, an inverse FFT was produced where only motile kinesin tracks remain. (C) The Directionality plugin (Jean-Yves Tinevez) was used, as described in the methods section, to deduct the frequency distribution of the directionality in the kymograph, reflecting kinesin velocities.

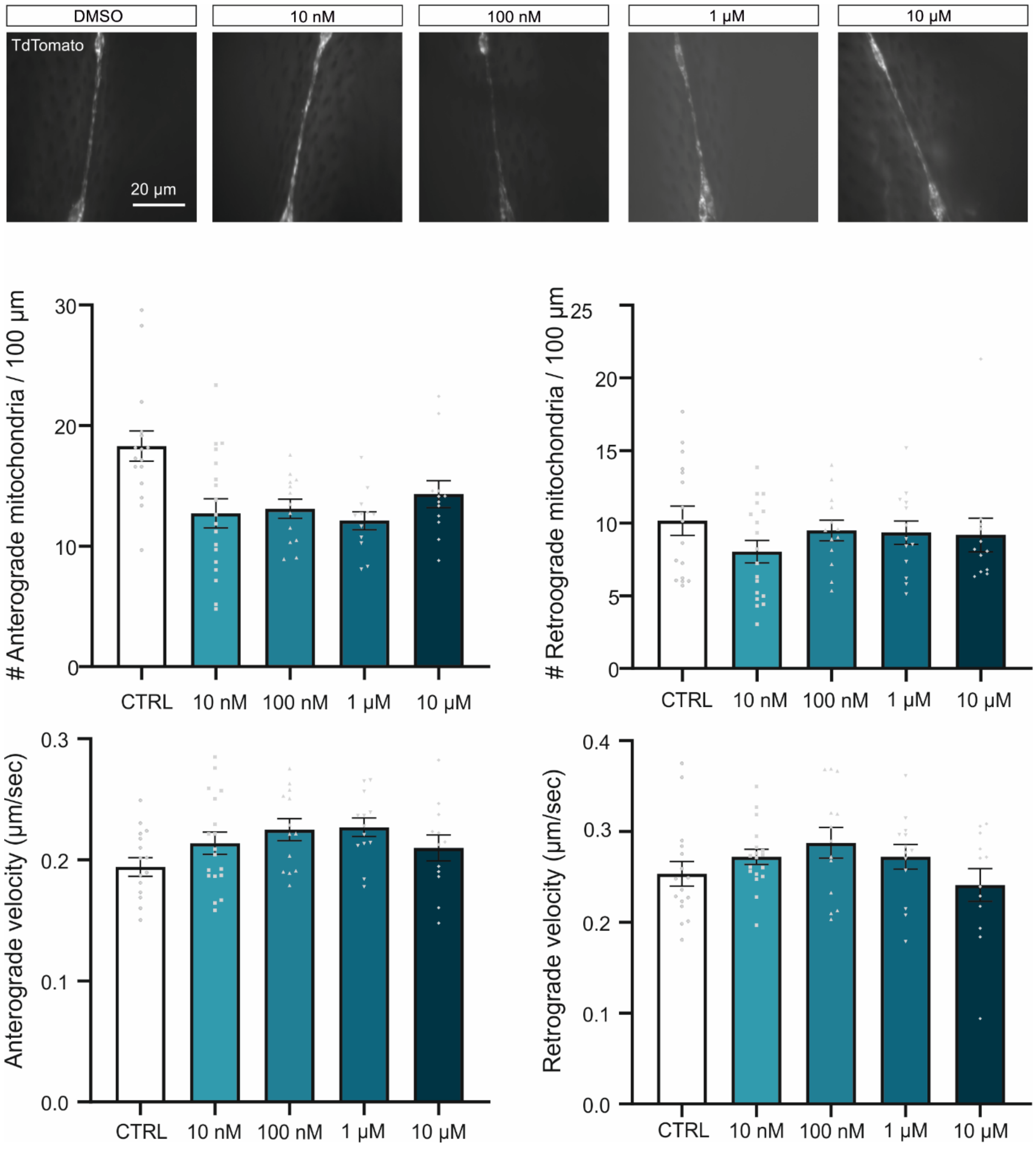


**Supplementary Figure 7. Pilot study for oral administration of taxol in Drosophila.**

Upper panels show sum projections of mitochondrial transport time lapses in axonal projections. Flies were treated with either DMSO as control or distinct concentrations of taxol (10 nM, 100 nM, 1 µM and 10 µM) for 24 hours before imaging. Graphs represent number of motile mitochondria in the antero- and retrograde direction and direction-specific velocities.
